# Exposure to the persistent organic pollutant 2,3,7,8-Tetrachlorodibenzo-p-dioxin (TCDD, dioxin) disrupts development of the zebrafish inner ear

**DOI:** 10.1101/2023.03.14.532434

**Authors:** Layra G. Cintrón-Rivera, Gabrielle Oulette, Aishwarya Prakki, Nicole M. Burns, Ratna Patel, Rachel Cyr, Jessica Plavicki

**Affiliations:** Department of Pathology and Laboratory Medicine, Brown University, 70 Ship St, Providence, RI, 02903

**Keywords:** zebrafish, dioxin, TCDD, ear, otic vesicle, semicircular canal

## Abstract

Dioxins are a class of highly toxic and persistent environmental pollutants that have been shown through epidemiological and laboratory-based studies to act as developmental teratogens. 2,3,7,8-tetrachlorodibenzo-p-dioxin (TCDD), the most potent dioxin congener, has a high affinity for the aryl hydrocarbon receptor (AHR), a ligand activated transcription factor. TCDD-induced AHR activation during development impairs nervous system, cardiac, and craniofacial development. Despite the robust phenotypes previously reported, the characterization of developmental malformations and our understanding of the molecular targets mediating TCDD-induced developmental toxicity remains limited. In zebrafish, TCDD-induced craniofacial malformations are produced, in part, by the downregulation of *SRY-box transcription factor 9b* (*sox9b*), a member of the SoxE gene family. *sox9b*, along with fellow SoxE gene family members *sox9a* and *sox10*, have important functions in the development of the otic placode, the otic vesicle, and, ultimately, the inner ear. Given that *sox9b* in a known target of TCDD and that transcriptional interactions exist among SoxE genes, we asked whether TCDD exposure impaired the development of the zebrafish auditory system, specifically the otic vesicle, which gives rise to the sensory components of the inner ear. Using immunohistochemistry, *in vivo* confocal imaging, and time-lapse microscopy, we assessed the impact of TCDD exposure on zebrafish otic vesicle development. We found exposure resulted in structural deficits, including incomplete pillar fusion and altered pillar topography, leading to defective semicircular canal development. The observed structural deficits were accompanied by reduced collagen type II expression in the ear. Together, our findings reveal the otic vesicle as a novel target of TCDD-induced toxicity, suggest that the function of multiple SoxE genes may be affected by TCDD exposure, and provide insight into how environmental contaminants contribute to congenital malformations.

**Highlights:**

1. The zebrafish ear is necessary to detect changes in motion, sound, and gravity.
2. Embryos exposed to TCDD lack structural components of the developing ear.
3. TCDD exposure impairs formation of the fusion plate and alters pillar topography.
4. The semicircular canals of the ear are required to detect changes in movement.
5. Following TCDD exposure embryos fail to establish semicircular canals.

## 1 Introduction

Exposure to environmental contamination is a persistent, global issue that shapes the health of aquatic species and humans alike. The complexity of exposures is increasing due to the generation of new contaminants produced from continuous industrial growth and inadequate environmental regulations, as well as the mobilization of legacy contamination by extreme weather events (Briggs, 2003). Aquatic environments are highly susceptible to the deleterious effects of toxic environmental contaminants with toxicant exposures in fish populations having the potential to produce morphological and behavioral changes that lead to decreased reproduction, health, and survival (Bashir et al., 2020; Melvin & Wilson, 2013)

Dioxins are a class of structurally related, widespread, toxic, and highly persistent contaminants released as byproducts of industrial practices such as fossil fuel combustion and medical waste incineration (Dopico & Gómez, 2015; Patrizi & de Cumis, 2018; Rysavy, Maaetoft-udsen, & Turner, 2013; White & Birnbaum, 2009). Dioxins are chemically stable teratogens that bio-persist and bio-accumulate in the food chain (Manzetti & Spoel, 2014). 2,3,7,8-tetrachlorodibenzo-p-dioxin (TCDD, dioxin) is the most potent and toxic dioxin congener (Manzetti & Spoel, 2014) with its toxicity mediated primarily through activation of the aryl hydrocarbon receptor (AHR), a ligand activated transcription factor (Carney, Prasch, Heideman, & Peterson, 2006; Kim et al., 2013; Mimura & Fujii-Kuriyama, 2003; Patrizi & de Cumis, 2018; Xu et al., 2015). Additional environmental pollutants, such as dioxin-like polychlorinated biphenyls (DL-PCBs) and polycyclic aromatic hydrocarbons (PAHs), also induce developmental toxicity through activation of AHR (Mimura & Fujii-Kuriyama, 2003; Sugden, Leonardo-Mendonça, Acuña-Castroviejo, & Siekmann, 2017).

In this study, we utilized the zebrafish (*Danio rerio*) model to identify and characterize novel targets of embryonic TCDD-induced developmental toxicity. Zebrafish are an excellent model for understanding the impact of environmental contamination on embryonic development, as they develop *ex vivo* and are optically transparent during early stages; thus, allowing for the visualization of morphological development in real time (Dai et al., 2014; Scholz et al., 2008; Yang et al., 2009). The rich repertoire of transgenic tools available in the zebrafish provide toxicologists with an opportunity to examine the adverse effects of contaminant exposure with a high degree of resolution. Previous research using zebrafish has shown that embryonic TCDD exposure disrupts the development of multiple organ systems including the nervous and cardiovascular systems and produces craniofacial malformations (Burns, Peterson, & Heideman, 2015; Hofsteen, Plavicki, Johnson, Peterson, & Heideman, 2013; Plavicki, Hofsteen, Peterson, & Heideman, 2013; Xiong, Peterson, & Heideman, 2008, and reviewed in Carney et al., 2006). Previous work demonstrated that inhibition of *sox9b*, a member of the SoxE gene family, is a critical molecular mediator of multiple TCDD-induced phenotypic endpoints. Given the critical role of multiple SoxE genes in teleost and mammalian ear development, we asked whether TCDD exposure affected the development of the auditory and vestibular systems, which are inherently intertwined(Barrionuevo et al., 2008; Liu et al., 2003; Park, Byung-Yong, 2010; Saint-Germain, Lee, Zhang, Sargent, & Saint-Jeannet, 2004; Szeto et al., 2022; Yan et al., 2005).

In zebrafish the otic vesicle, also referred to as the ear, is required to sense changes in sound, gravity, and motion (Abbas & Whitfield, 2010; Baxendale & Whitfield, 2016; Dutton et al., 2009; Geng et al., 2013; Whitfield, 2002; Whitfield, Riley, Chiang, & Phillips, 2002). Fish depend on sensory cues to acquire nutrients, escape from prey, and survive in changing currents. Therefore, proper ear development is critical for fish health and fitness. The zebrafish ear is composed of three semicircular canals, three cristae, two otoliths, and two macula (Bever & Fekete, 2002; Liu et al., 2003; Whitfield, 2020; Whitfield et al., 2002). Together, the dorsolateral septum, anterior pillar, ventral pillar, and posterior pillar establish three semicircular canals required to detect changes in angular acceleration and rotational movement (Baxendale & Whitfield, 2016). The cristae, otoliths, and macula house sensory hair cells, which are necessary for vestibular function. Failure to establish semicircular canals or to develop the hair cells of the ear results in sensory perception defects.

To examine how early embryonic exposure to TCDD affects otic development, we exposed zebrafish embryos carrying a transgenic reporter for *sox10*, an early neural crest marker required for development of the otic vesicle epithelium as well as the peripheral nervous system, melanocytes, heart, and jaw (Dutton et al., 2009). Embryos were exposed to a 1-hour waterborne exposure of 10 parts-per-billion (ppb) TCDD at 4 hpf, an exposure paradigm that results in an internal body burden of 14.7 ±2.04 pg/embryo at 48 hpf and 9.78 ±1.31 pg/embryo at 72 hpf respectively (Kossack, Manz, Martin, Pennell, & Plavicki, 2023). Using confocal microscopy of fixed and live samples, we characterized the effects of exposure on otic vesicle morphology, quantifying fusion plate formation and pillar topography. We determined that TCDD exposure significantly disrupted the structural development of the ear, resulting in the failure of semicircular canals to form. In addition to the significant morphological malformations observed, we found TCDD exposure reduces collagen type II expression in the developing otic vesicle. Our data demonstrate that the developing otic vesicle is a target of TCDD toxicity and, given our prior work demonstrating TCDD exposure disrupts olfactory system development, lead us to conclude that embryonic TCDD exposure impairs development of multiple sensory modalities.

## 2 Methods

### 2.1 Zebrafish Husbandry and Transgenic Lines

All experiments in this study were conducted utilizing zebrafish from the Plavicki Lab Zebrafish Facility. All procedures were covered by protocols approved by the Institutional Animal Care and Use Committee (IACUC) at Brown University in adherence with the National Institute of Health’s “Guide for the Care and Use of Laboratory Animals.” Transgenic zebrafish lines were kept in an aquatic housing system (Aquaneering Inc., San Diego, CA) with automatic pH and conductivity stabilization, centralized filtration, reverse osmosis (RO) water purification, temperature maintenance (28.5 ± 2°C), and ultraviolet (UV) irradiation for pathogen control. Fish were reared and maintained with a 14 hour:10 hour light:dark cycle, according to Westerfield (2000). The Plavicki Lab Zebrafish Facility undergoes routine monitoring for disease including the semiannual quantified Polymerase Chain Reaction (qPCR) panels to detect common fish pathogens.

Adult zebrafish were spawned in 1.7L slope tanks (Techniplast, USA). Fish were placed in tanks 15-18 hours prior to breeding with a transparent divider that separated males and females. The following day the divider was removed within 2-3 hours of the initiation of the light cycle. Embryos were collected over the course of 1 hour and subsequently reared in fresh egg water (60 mg/l Instant Ocean Sea Salts; Aquarium Systems, Mentor, OH) in 100 mm non-treated culture petri dishes (CytoOne, Cat. No. CC7672-3394). Embryonic and larval zebrafish were maintained in temperature regulated incubators with a 14 hour:10 hour light:dark cycle (Powers Scientific Inc., Pipersville, PA). At 24 hpf, embryos were manually dechorionated using Dumont #5XL forceps (Cat. No. 11253-10, Fine Science Tools, Foster City, CA) and moved to fresh egg water containing 0.003 % 1-phenyl-2-thiourea (PTU, Sigma) to prevent pigment deposition. The following transgenic lines were used in this study: *Tg*(*Sox10:RFP*)(Kirby et al., 2006) and *TgBAC*(*cyp1a:NLS:EGFP*) (Kim et al., 2013).

### 2.2 TCDD exposure

Embryonic TCDD and dimethyl sulfoxide (DMSO) control exposures were performed in accordance with Plavicki et al., (2013) (Plavicki et al., 2013). Within 1 hour of adults spawning, embryos were collected and maintained in a temperature-controlled incubator for future assessment. At 3 hpf fertilized embryos were evaluated and selected based on healthy developmental quality. Selected embryos were exposed at 4 hpf to either a 0.1% DMSO vehicle control or to a 10 ppb TCDD dosing solution for 1 hour while gently rocking. Twenty embryos were exposed per 2 mL of dosing solution in 2 mL glass amber vials (Cat. No. 5182-0558, Agilent, Santa Clara, CA). Control and dosing solutions were made by adding solution constituents to amber vials already containing 20 embryos and gently rocking. For TCDD exposures, a 10 ppb TCDD (ED-901-B, Cerriliant, St. Louis, MO) dosing solution was prepared by adding 2 mL of egg water and 2 μL of pre-prepared TCDD stock (10ng/mL). Vehicle controls were exposed to a 0.1% DMSO control dosing solution, which is matched to the DMSO concentration in the TCDD dosing solution. Amber vials and caps were wrapped with Parafilm^®^ to limit evaporation.

### 2.3 Immunohistochemistry

Zebrafish embryos and larvae were collected at 72 hpf and fixed in groups of 10-15 fish per 1.5 mL Eppendorf^®^ Safe-Lock microcentrifuge tube (Millipore Sigma, Cat. No. T9661-500EA) in 4% paraformaldehyde (PFA, Sigma-Aldrich, Cat. No. P6148) overnight at 4°C while gently rocking. Samples were washed 5 times with PBS-T (phosphate buffered solution + 0.6% Triton-X 100) post-fixation. Samples were permeabilized with acetone for 5 minutes at −20°C. Post-permeabilization, samples were placed in blocking solution (PBS-T + 4% bovine serum albumin, BSA) overnight while gently rocking at 4°C. After blocking, samples were incubated with 1° antibodies at 4°C for 1 day. Samples next underwent 5 quick washes with PBS-T and then washed overnight in fresh PBS-T at 4°C. Once PBS-T wash solution was removed, the 2° antibody and Hoechst stain (1:10,000) were added to the samples and left to incubate for approximately 18 hours at 4°C. 2° antibody and Hoechst were removed by 5 quick washes performed with PBS-T followed by overnight wash at 4°C in PBS-T. PBS-T solution was fully removed from the Eppendorf tubes using a Samco™ Fine Tip Transfer Pipette (Millipore Sigma, Cat. No. Z350605-500EA) and VECTASHIELD Mounting Media (Vector Labs, Cat. No. H-1000) was added for clearing and storage. Samples were stored for a minimum of 24 hours in VECTASHIELD prior to imaging. The 1° antibody used was Collagen Type II (II-II6B3) (1:100, mouse monoclonal [IgG1]). II-II6B3 was deposited to the Developmental Studies Hybridoma Bank (DSHB) by Linsenmayer, T.F. (DSHB Hybridoma Product II-II6B3). The 2° antibody used was Alexa Fluor^®^ 488 Goat antiMouse IgG1 Cross-Adsorbed Secondary Antibody (1:100,ThermoFisher Scientific, Cat. A21121)

### 2.4 Microscopy and Image Analysis

Live embryos and larvae were anesthetized with 0.02% tricaine (MS-222) for embedding and subsequent imaging. Both live and fixed embryos were mounted in 2% low-melting agarose (Fisher Scientific, bp1360-100) in 35 mm glass bottom microwell dishes (MatTek, Part No. P35G-1.5-14-C). Confocal imaging was performed using a Zeiss LSM 880 confocal with Airyscan Fast. Z-stacks spanned approximately 110 μm with images collected at 0.69 μm intervals. Images were analyzed and prepared for publication using Adobe Photoshop, Adobe Illustrator, Zeiss Zen Black, and Zeiss Zen Blue software. Graphs were generated for data visualization utilizing GraphPad Prism and R studio.

### 2.5 Statistical Analysis

Chi-squared and Fishers exact test were used to assess differences between categorical data collected from the control and TCDD-exposed groups. Significant differences between groups were determined using Chi-squared and Fishers exact test with a 95% confidence interval in GraphPad Prism. Significance is denoted as follows: *p < 0.05, **p < 0.01, ***p < 0.001, ****p <0.0001.

## 3 Results

### 3.1 Cyp1a, a biomarker of AHR activation, is induced in the developing otic vesicle following TCDD exposure

TCDD-induced AHR activation induces expression of cytochrome P450 1 A (*cyp1a*), a xenobiotic metabolizing enzyme and a biomarker of AHR activation (Carney et al., 2006; Kim et al., 2013; Mimura & Fujii-Kuriyama, 2003; Xu et al., 2015). Correspondingly, *cyp1a* mRNA or protein expression can be used to identify cellular targets of TCDD-induced developmental toxicity. To determine if the developing otic vesicle was a target of TCDD toxicity, we exposed transgenic zebrafish embryos carrying a green fluorescent protein reporter in the start codon of Cyp1a (*Tg*(*cyp1a:nls-egfp*)) to 10 ppb TCDD at 4 hpf and used confocal microscopy to visualize Cyp1a expression at 48 and 72 hpf. Previous work has shown that during early stages of development there is endogenous expression of Cyp1a in the ear (Kim et al., 2013). Therefore, we sought to determine if TCDD exposure led to an expansion of the Cyp1a expression domain in the developing otic vesicle. Our findings show that exposure to TCDD results in robust expansion of the Cyp1a expression domain in the ear at both 48 and 72 hpf (Fig. 2 & Fig. 3, Supp. Fig 1).

**Figure 1.**
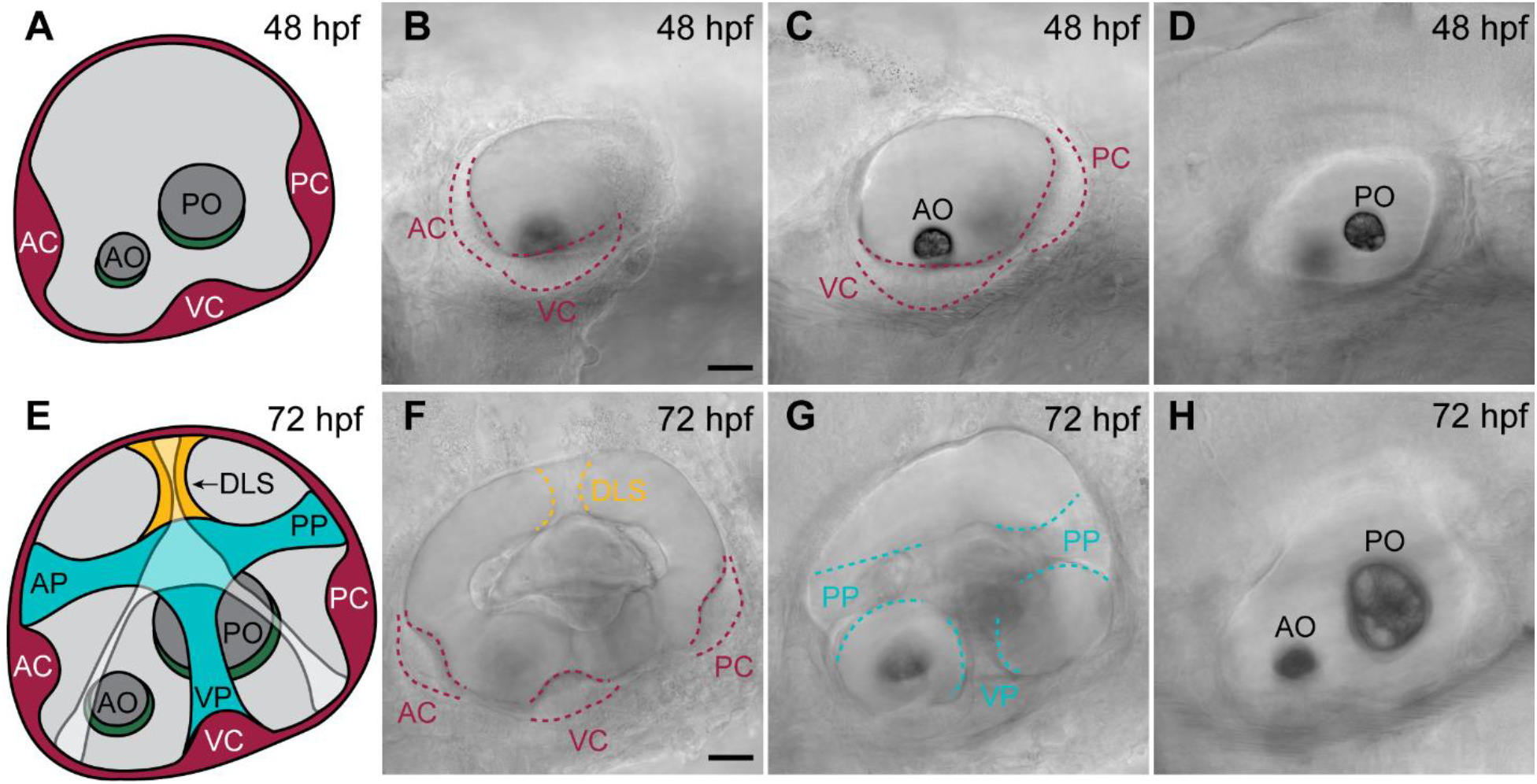
Anatomy of the wild-type zebrafish ear. (A, E) Schematics of wild-type zebrafish ear at 48 hpf (A) and 72 hpf (E). (B-D) Representative brightfield images of live wild-type ears showing the developing cristae and otoliths at 48 hpf. (F-H) Representative brightfield images of live wild-type ears capturing the cristae, otoliths, pillars, and dorsolateral septum at 72 hpf. Abbreviations: anterior crista (AC), ventral crista (VC), posterior crista (PC), anterior pillar (AP), ventral pillar (VP), posterior pillar (PP), dorsolateral septum (DLS), anterior otolith (AO), posterior otolith (PO). All images are 40x. Scale bar in B & F is 25μm.

**Figure 2.**
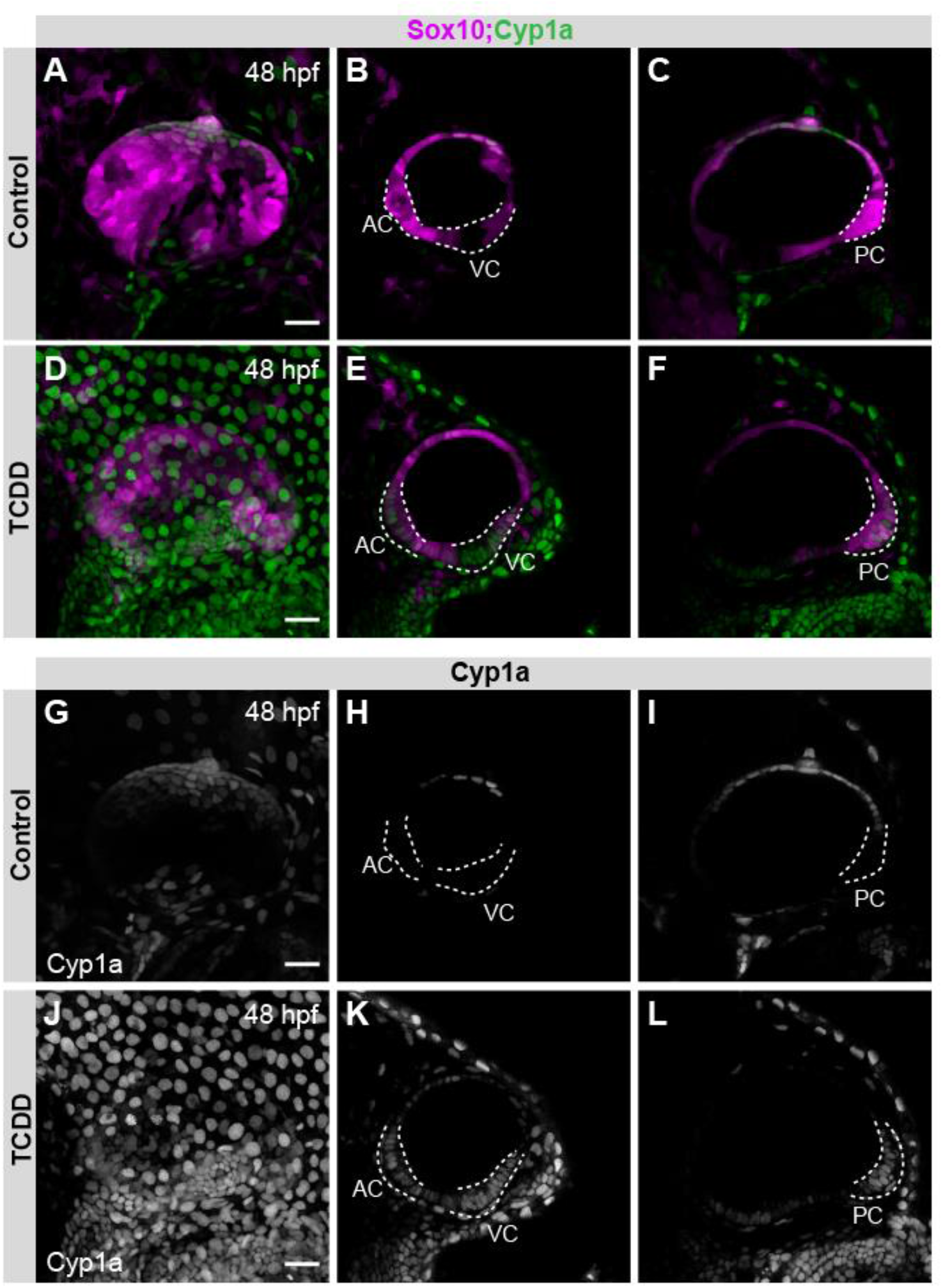
TCDD exposure expands the Cyp1a expression domain in the developing embryonic ear. (A-F) Confocal micrographs of control (A-C) and TCDD-exposed (D-F) embryos carrying reporters for *sox10* (magenta) and *cyp1a* (green) *[Tg(sox10:RFP); TgBAC(cyp1a:NLS:EGFP)]. sox10* expression encompasses the otic epithelium and *cyp1a* acts as a biomarker of AHR activation. (H-I, K-L) Representative slices taken from confocal z-series capturing *cyp1a* induction (white) in control (G-I) and TCDD-exposed (J-L) embryos. Maximum intensity projections of confocal z-series are shown in A, D, G, & J. Abbreviations: anterior crista (AC), ventral crista (VC), and posterior crista (PC). Scale bar in A, D, G, & J = 25μm. n=8-10/group with 3 a minimum of replicates.

**Figure 3.**
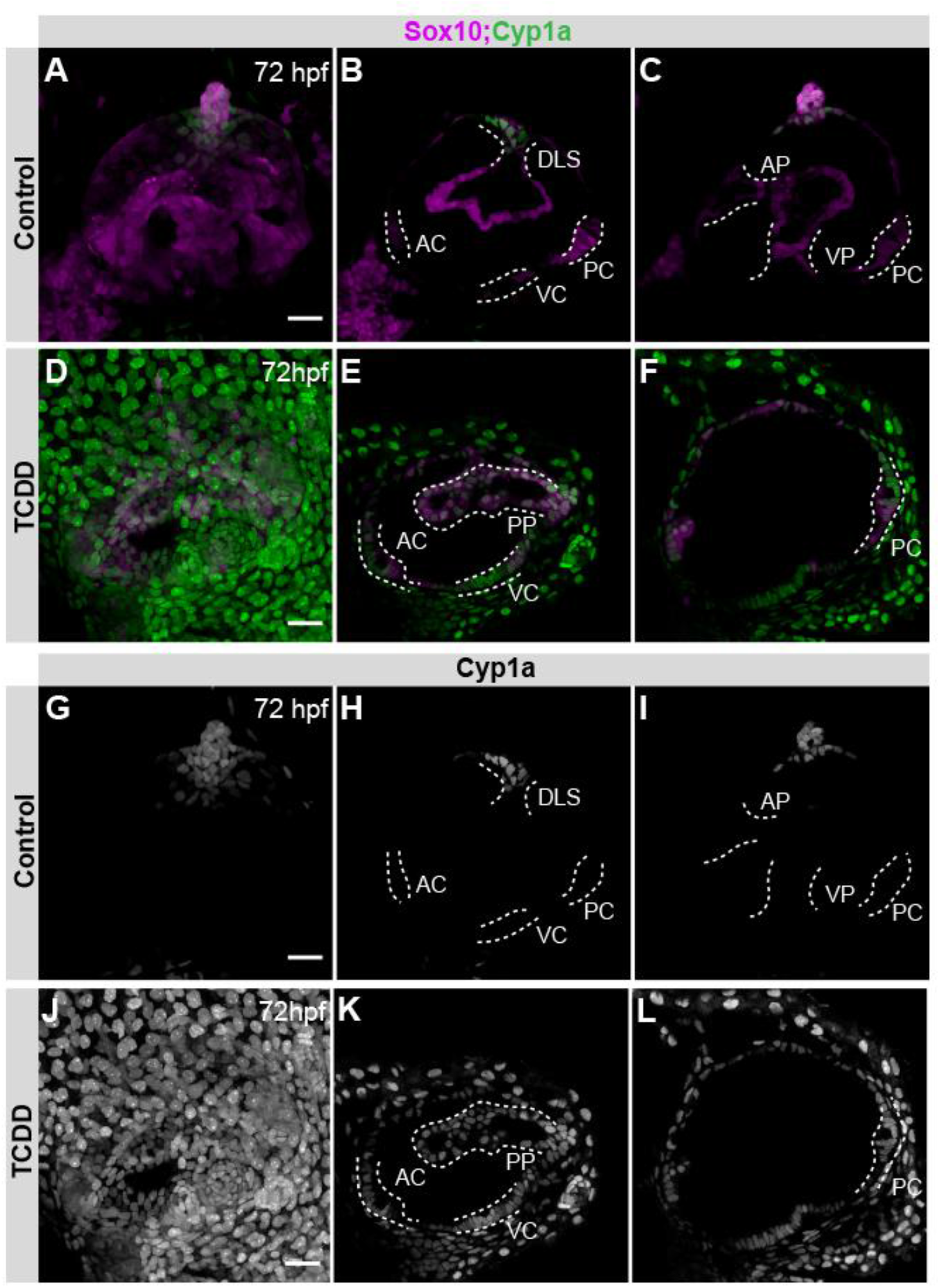
TCDD-induced expansion of Cyp1a expression is maintained in the larval ear. (A-F) Confocal micrographs of control (A-C) and TCDD-exposed (D-F) larvae carrying reporters for *sox10* (magenta) and *cyp1a* (green) *[Tg(sox10:RFP); TgBAC(cyp1a:NLS:EGFP)]*. (G-L) Cyp1a transgenic reporter in black and white in controls (G-I) and TCDD exposed (J-L) fish. Maximum intensity projections of confocal z-series are shown in A, D, G, & J. Abbreviations: anterior crista (AC), ventral crista (VC), posterior crista (PC), anterior pillar (AP), ventral pillar (VP), posterior pillar (PP), dorsolateral septum (DLS). Scale bar in A, D, G & J is 25μm. n=10-13/group with a minimum of 3 replicates.

By 48 hpf, the otic vesicle has developed Sox10+ otic epithelium throughout the ear. At this developmental stage, the three cristae have been established; however, the pillar structures are not yet present (Fig. 1 A-D). Our data showed that by 48 hpf control embryos had Cyp1a expression in the dorsal most portion of the ear, which we will refer to as the apex of the ear, and within a small subset of cells within in the anterior crista. (Fig. 2 A-C, G-I; Supp. Fig. 1). In contrast, in TCDD-exposed embryos, Cyp1a was expressed throughout the anterior crista, ventral crista, and posterior crista at 48 hpf, suggesting the cristae are an early target of TCDD-induced toxicity (Fig. 2 D-F, J-L; Supp. Fig 1).

By 72 hpf, the ear has developed four additional structures: the anterior pillar, ventral pillar, posterior pillar, and dorsolateral septum, which together establish the semicircular canals required for vestibular function (Fig. 1 E-H). At 72 hpf, only the apex and the dorsolateral septum show expression of Cyp1a in control larvae (Fig. 3 A-C, G-I; Supp. Fig. 1). In contrast, the Cyp1a expression domain in TCDD exposed larvae at 72 hpf encompassed the apex, anterior crista, ventral crista, posterior crista, and the present anterior pillar, ventral pillar, posterior pillar, and dorsolateral septum (Fig. 3 D-F, J-L; Supp. Fig. 1). Thus, suggesting that the otic vesicle cristae and the pillar structures that make up the semicircular canals continue to be targets of TCDD-induced toxicity.

The three cristae house sensory hair cells that aid in, but are not required for, accurate tethering of the otoliths (Whitfield, 2020; Whitfield et al., 2002). Since TCDD induced Cyp1a expression in the cristae, we evaluated ears for the presence of otoliths and found that the otoliths were present in all 48 hpf control and TCDD exposed embryos (n=11/group with 3 replicates); as well as in all 72 hpf control and TCDD exposed embryos (n=8-9/group with 3 replicates).

### 3.2 Ears of TCDD exposed embryos display structural deficits

Proper establishment of the semicircular canals requires the development and subsequent fusion of the anterior pillar, ventral pillar, posterior pillar, and dorsolateral septum. Given that the expanded pattern of Cyp1a expression at 72 hpf encompassed the developing pillars, we sought to determine if TCDD exposure disrupts development of the semicircular canals. Semicircular canals form between 60 hpf to 90 hpf (Baxendale & Whitfield, 2016). Therefore, to determine if TCDD exposure disrupted development of the semicircular canals, we characterized pillar fusion at 72 hpf and 96 hpf. We began by quantifying the presence of structural components of the ear and saw that by 72 hpf, 95% of controls had all of the semicircular structural components present. Meanwhile, only 4.17% of TCDD exposed embryos showed the presence of all structures (Fig. 4A). 75% of TCDD embryos lacked the ventral pillar, 20.83% the anterior pillar, and 8.33% the posterior pillar (Fig. 4B). Only 5% of control embryos were missing the ventral pillar, which likely reflected variability in developmental timing of ventral pillar development (Fig. 4B). In contrast, TCDD exposed fish showed a range of deficits with 70.83% of embryos lacking at least two structural components and 25% lacking one structure alone (Fig. 4A). Coincidence of a structural absence varied in severity with 8.33% of TCDD exposed fish having no structural components present and 12.50% showing coincident deficits in anterior pillar, ventral pillar, and dorsolateral septum establishment (Fig. 4A). The most common coincident phenotype following TCDD exposure was absence of both the ventral pillar and dorsolateral septum, 50% of embryos presented this defect (Fig. 4A). The dorsolateral septum was the most frequently absent structure with 91.67% of TCDD exposed embryos showing a deficit (Fig. 4B).

**Figure 4.**
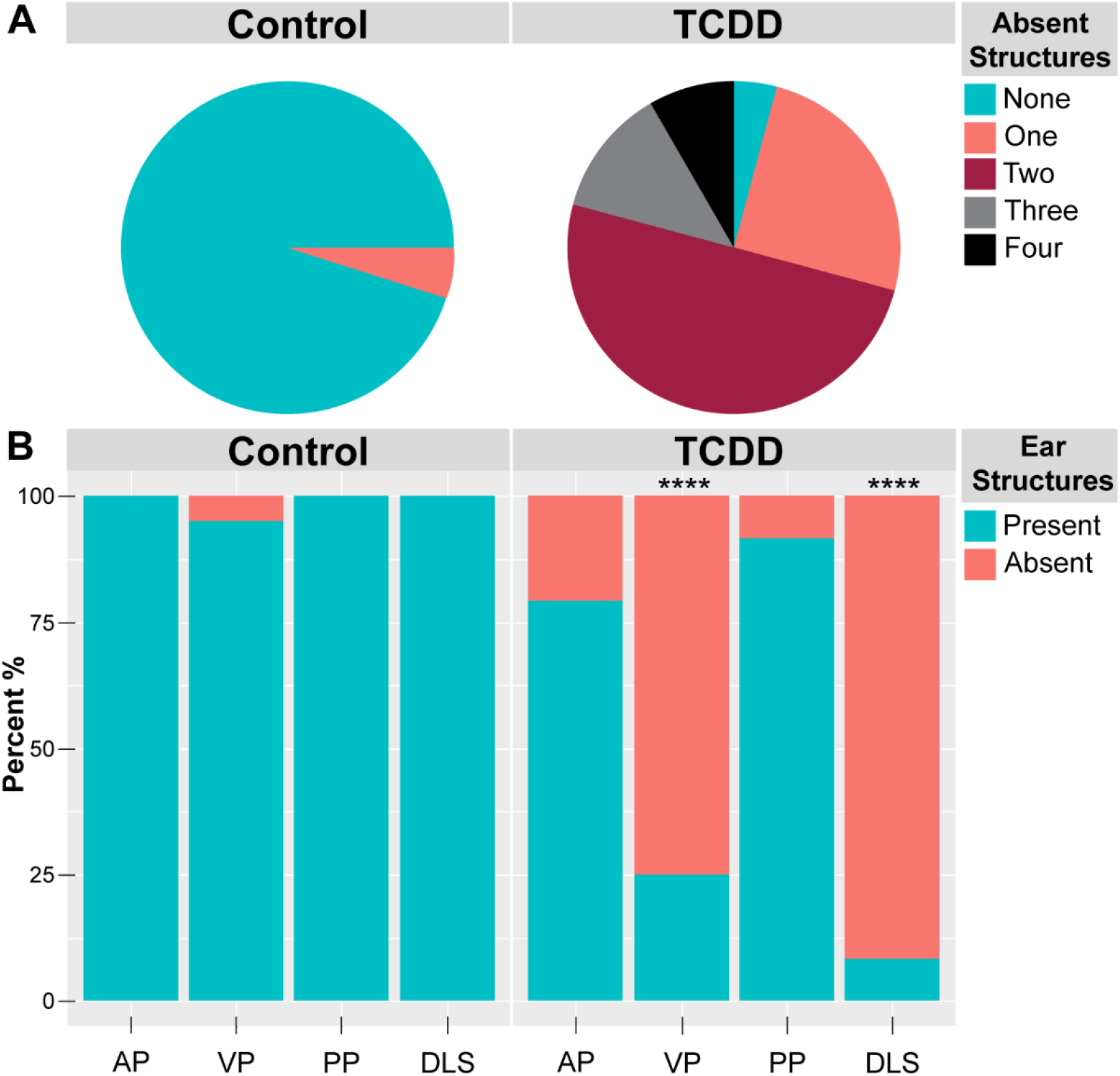
Quantification of structural deficits induced by TCDD exposure. (A) Percentage of coincident absent structures in control and TCDD exposed larval at 72 hpf. (B) Percentage of specific ear structure absence. Abbreviations: anterior pillar (AP), ventral pillar (VP), posterior pillar (PP), dorsolateral septum (DLS). n=20-24/group with a minimum of 3 replicates. ****p <0.0001.

The rates of coincident phenotypes varied at 96 hpf compared to those at 72 hpf. At 96 hpf the percentage of fish with all structures present increased to 100% in the control group and 15% in the TCDD exposed group, compared to 95% and 4.17% respectively at 72 hpf. Additionally, at 96 hpf, only 60% of TCDD exposed embryos, compared to 70.83% at 72 hpf, had at least two coincident phenotypes. However, while only 8.33% of TCDD exposed larvae showed absence of all structural components at 72 hpf, this increased to 35% at 96 hpf (Supp. Fig. 2). Overall, these data indicate that TCDD exposure leads to significant structural deficits with ranging degrees of severity at both 72 hpf and 96 hpf.

### 3.3 Embryos dosed with TCDD fail to establish a fusion plate

In addition to pillar development, complete pillar fusion is required for establishment of semicircular canals. For pillar fusion to occur the anterior pillar, ventral pillar, and posterior pillar must be present and connect at the fusion plate in the center of the ear (Whitfield et al., 2002). To capture whether complete fusion occurred, we scored larvae for complete and incomplete pillar fusion at both 72 hpf and 96 hpf. Ears were classified as having an incomplete fusion phenotype if the anterior pillar, ventral pillar, and posterior pillar did not form a fusion plate. Therefore, embryos that lacked a pillar or had pillars present that did not fuse were categorized as having incomplete fusion. In controls, 30% of larvae showed an incomplete fusion of the ventral pillar at 72 hpf (Fig. 5). This was likely due to variability in development, since at 96 hpf 0% of controls were scored as having incomplete fusion phenotypes. In contrast, 100% TCDD exposed embryos showed a defect in fusion at both 72 hpf and 96 hpf (Fig. 5 and Supp. Fig. 3).

**Figure 5.**
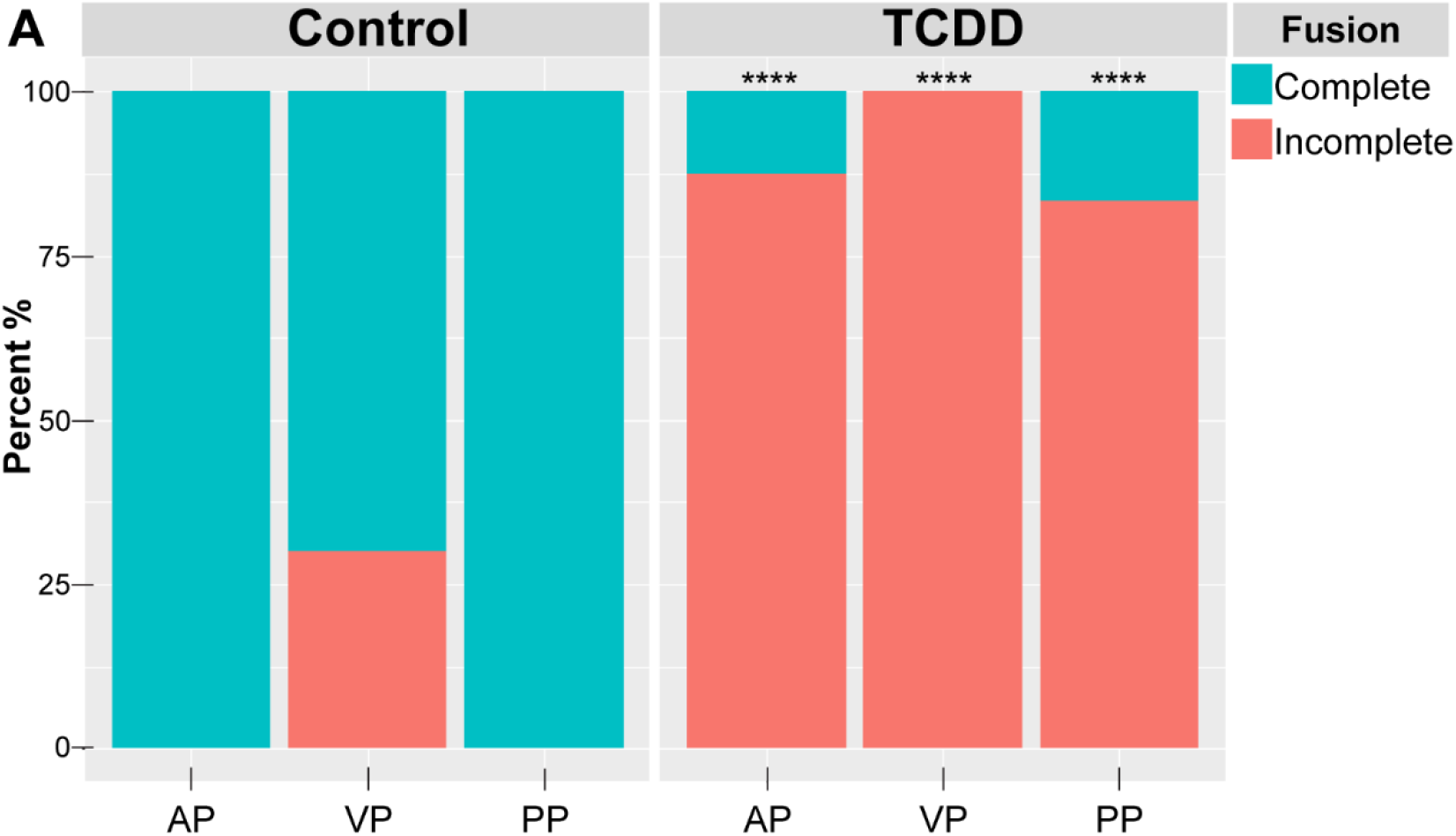
Quantification of incomplete pillar fusion. Percent of structures that failed to establish a connection at fusion plate in control and TCDD exposed embryos at 72hpf (A). Abbreviations: anterior pillar (AP), ventral pillar (VP), posterior pillar (PP). n=20-24/group with a minimum of 3 replicates. ****p < 0.0001.

Incomplete fusion phenotypes can occur even if all pillars are present. A pillar was categorized as having incomplete fusion if it was absent or if it did not fuse with at least one of the other pillars. At 72 hpf, 100% of TCDD exposed embryos displayed incomplete fusion of the ventral pillar; 75% of embryos lacked a ventral pillar all together, while 25% had a ventral pillar present that did not link at the fusion plate. Likewise, 87.5% of larvae previously dosed with TCDD showed incomplete fusion of the anterior pillar by 72 hpf; 20.83% of larvae showed absence of the anterior pillar, whereas 66.67% had an existing anterior pillar that did not reach the fusion plate. Finally, 83.33% of TCDD exposed larvae demonstrated an incomplete posterior pillar fusion phenotype; 8.33% of larvae were missing the posterior pillar, whereas 75% lacked adequate formation of the fusion plate formation (Fig. 5). Although we have delineated fusion defects per pillar by categorizing which pillars did not fuse at the plate, it is important to note that presence of all three pillars is required to form the fusion plate. Therefore, after exposure to TCDD 100% embryos at 72 hpf fail to establish the fusion plate. We were able to capture these defects in establishment and fusion of pillars occurring between 52-72 hpf though time-lapse confocal microscopy (Supp. Movie 1).

Lastly, to determine if the pillars in TCDD exposed fish fuse at a later time point, we assessed fusion plate establishment at 96 hpf. Consistent with data at 72 hpf, 100% of TCDD exposed larvae had incomplete pillar fusion at 96 hpf. 85% showed defects in the anterior pillar’s ability to reach and fuse with the fusion plate, 95% showed defect at the ventral pillar, and 95% showed defect at the posterior pillar (Supp. Fig. 3). Together, these data indicate that embryonic exposure to TCDD impairs establishment of the semicircular canals by altering both establishment and fusion of pillar structures.

### 3.4 TCDD exposure changes topography of ear structures

Through the process of quantifying semicircular canal establishment, we noted all pillars in control larvae develop in a stereotyped position relative to the adjacent crista. In control fish, the anterior pillar buds above the anterior crista, the ventral pillar buds on top of or slightly to the right of the ventral crista, and the posterior pillar buds above the posterior crista. In contrast, the topography of pillars in TCDD exposed fish was inconsistent; therefore, we quantified how many fish showed a topographical displacement of pillar development. Pillars that were categorized as displaced were present and budding, but in an incorrect position relative to the pillar-crista relationship observed in all control larvae. 100% of the pillars in control larvae develop in the same topographical relation to the cristae. At 72 hpf, 59.09% TCDD exposed fish showed displacement of the posterior pillar and 9.09% showed a coincident displacement of both the anterior and posterior pillars (Fig. 6A). Specifically, the posterior pillar developed under the posterior crista, while the anterior pillar developed on top of or under the anterior crista (Fig. 6B-G). Meanwhile, at 96 hpf, 83.33% of TCDD exposed embryos had one pillar displaced, but no coincident displacements (Supp. Fig. 4). This data indicates that TCDD exposure altered the patterning and overall structural development of the otic vesicle.

**Figure 6.**
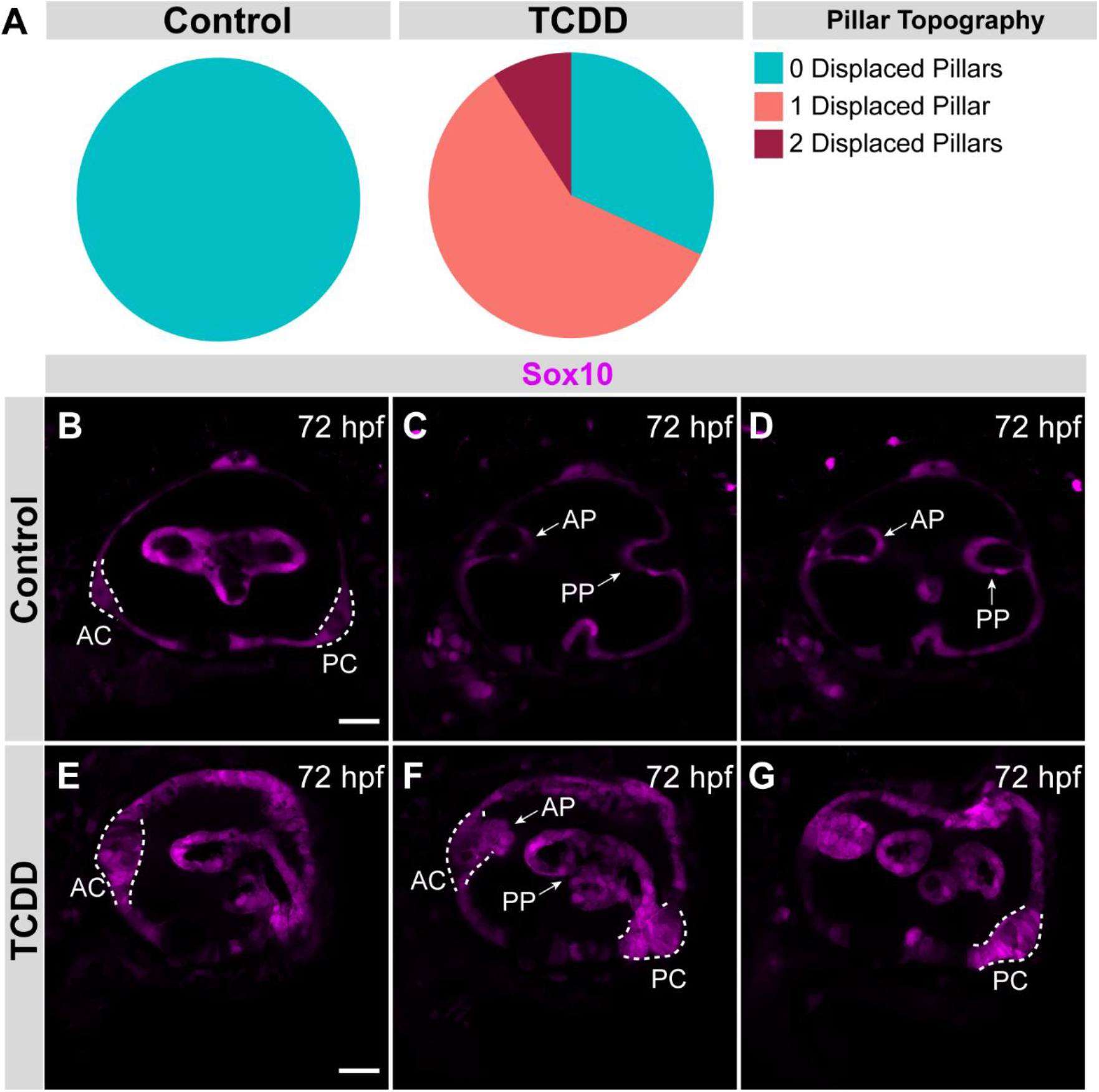
Pillar topography defect quantification and phenotypes. Percent of control and TCDD exposed embryos with zero, one, or two displaced pillars at 72 hpf (A). Confocal slices capturing the fluorescent reporter *Tg(Sox10:RFP*), in magenta, in the developing otic epithelium at 72hpf with a 40x magnification (B-G). Control ear displaying typical pillar topography (B-D). TCDD exposed ear exhibiting displacement of both the anterior and posterior pillar (E-G). Abbreviations: anterior pillar (AP), ventral pillar (VP), posterior pillar (PP), dorsolateral septum (DLS), anterior crista (AC), posterior crista (PC). 25μm scale bar for reference (B,E). n=20-24/group with a minimum of 3 replicates.

### 3.5 The collagen type II expression in the ear is reduced following TCDD exposure

Collagen type II (Col2a1) is expressed early on in the developing otic vesicle epithelium, pillar structures, and in the anterior macula (Chung et al., n.d.; Dale & Topczewski, 2011; Williams, 2022; Yan et al., 2005). Col2a1 plays an important role in maintaining structural integrity in developing skeletal structures. Given that *Sox9*, a target of AHR signaling in mammals, is a direct regulator of Col2a1 (Bell et al., 1997) and that *sox9b*, one of 2 orthologues of *Sox9*, is known to be repressed in multiple zebrafish tissues following TCDD exposure, we asked if expression of Col2a1 in the ear was affected by embryonic TCDD exposure. By 72 hpf, all control embryos showed expression of Col2a1 in the otic epithelium and present pillar structures. Meanwhile, 90% of TCDD exposed fish had reduced Col2a1 expression in the ear (Fig. 7).

**Figure 7.**
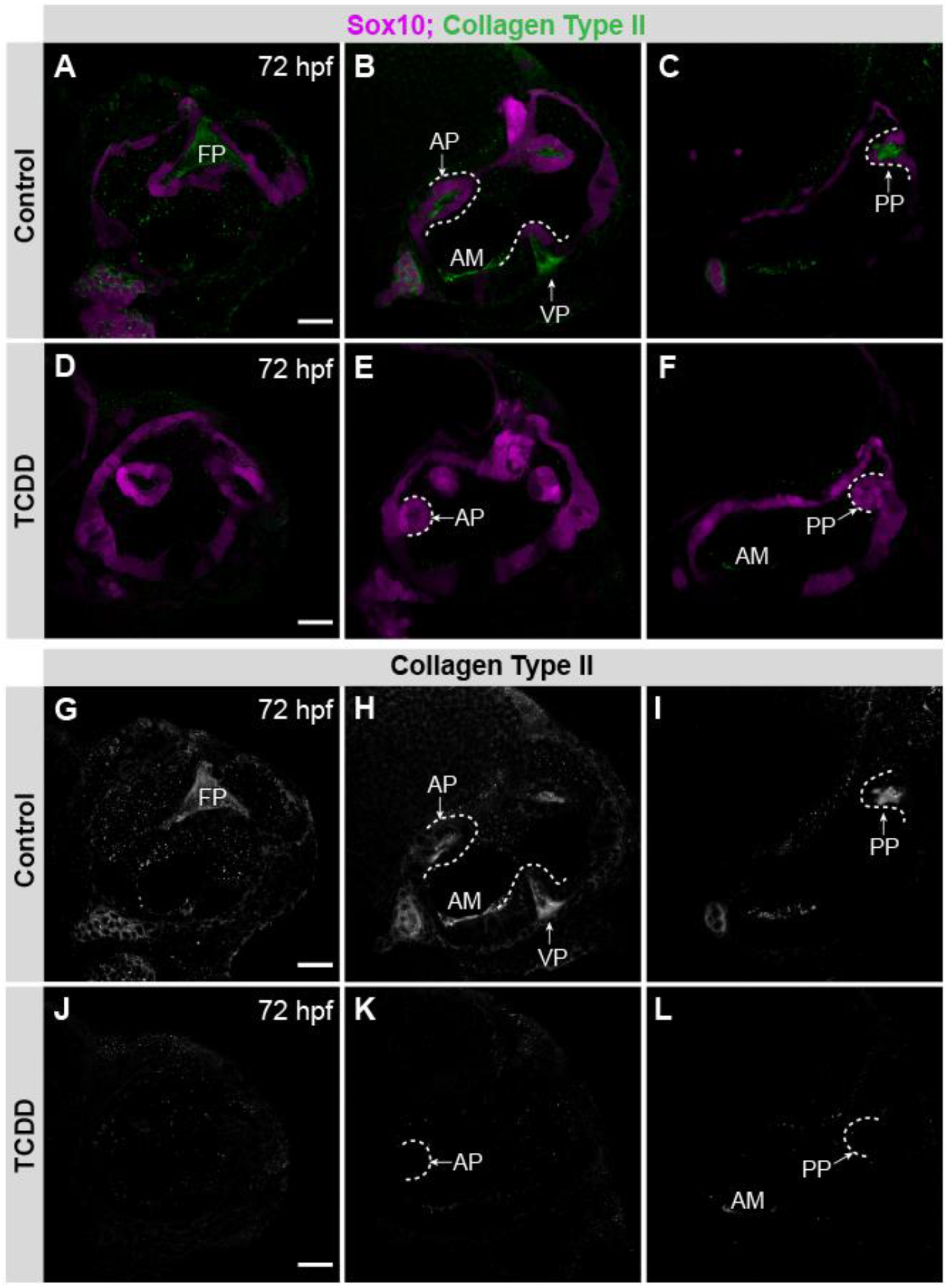
Loss of collagen type II expression in the ear following TCDD exposure. (A-L) 40x confocal micrographs highlighting different portions of the developing ear at 72 hpf. Control (A-C) and TCDD exposed embryos (D-F) expressing the fluorescent reporter *Tg*(*sox10:RFP*) (magenta) and immunostained for collagen type II (green). (G-L) Black and white images of collagen type II in control (D-F) and TCDD exposed (J-L) larval ears. Abbreviations: fusion plate (FP), anterior macula (AM), anterior pillar (AP), ventral pillar (VP), posterior pillar (PP). Scale bar in A,D,G,J = 25μm.n=11-8/group with 3 replicates.

## 4 Discussion

### 4.1 TCDD exposure disrupts sensory system development

Organisms rely on their sensory modalities for the acquisition of nutrients, mating, and evasion of predators. Therefore, the proper structural development of sensory organs as well as appropriate neural innervation of these organs is essential for survival and fitness. Zebrafish possess all of the primary sensory modalities: hearing, balance, taste, touch, smell, and vision (Moorman, 2001). Research in fish and mammals has demonstrated that TCDD exposure at different life stages results in neurodevelopmental phenotypes in the CNS including impaired neurogenesis and disrupted development of the olfactory bulb, cerebellum, optic tectum, habenula, and hippocampus (Collins et al., 2008; Hill, Howard, Strahle, & Cossins, 2003; Iida, Kim, Murakami, Shima, & Iwata, 2013; Kimura et al., 2016; Williamson, Gasiewicz, & Opanashuk, 2005). However, considerably less is known about how TCDD exposure effects sensory system development. In this study, we identify the developing zebrafish ear as a novel target of TCDD-induced toxicity, revealing that TCDD exposure impairs development, fusion, and topography of essential ear structures. The ear is required for sensing changes in sound and, in conjunction with the lateral line, detecting changes in gravity, movement, and rotation. TCDD-induced disruption of ear development, therefore, has the potential to impair a number of ecologically relevant behaviors. While we did not observe gross malformations in the neural innervation of the ear, more nuanced studies using genetically encoded calcium indicators and/or electrophysiology could reveal neurofunctional changes in the vestibuloacoustic nerve that innervates the hair cells and/or the octavolateralis efferent system that carries information from the auditory vestibular system to the periphery (Beiriger, Narayan, Singh, & Prince, 2021; Blaxter, 1987). Previous work examining the impact of TCDD exposure on peripheral nervous system development in the red seabream found multiple cranial nerves to be disrupted by TCDD exposure; however, these studies did not specific examine the effects of TCDD exposure on the vestibuloacoustic nerve (Iida et al., 2013).

Previous work conducted in our lab showed that early TCDD exposure disrupts development of the olfactory organs and olfactory bulb, the structures required to detecting and coding olfactory cues (Martin et al., 2022). Additionally, while it is beyond the scope of this paper, we note that TCDD-exposed embryos and larvae examined in this study had altered eye morphology, indicating that the eye is also a target of TCDD-induced toxicity (Supp. Fig. 5). Consistent with our observation that TCDD-induced dysregulation of AHR signaling alters eye development, the invertebrate homologs of *AHR, spineless*, is necessary for retinal development (Wernet et al., 2006). Together, these studies suggest TCDD toxicity impairs development of multiple sensory systems.

### 4.2 Understanding the molecular mechanisms that mediate TCDD-induced otic phenotypes

Our study characterized the effects of TCDD toxicity on otic vesicle development. To our knowledge, no previous studies have described the ototoxic effects of TCDD in zebrafish or other aquatic animals. Zebrafish mutant analysis studies have revealed key signaling pathways and transcriptional regulators of otic vesicle development and semicircular canal establishment (Whitfield et al., 2002). These mutants highlight potential targets of interest in the quest to uncover the molecular mechanisms of TCDD-induced toxicity. Mutants with defects in semicircular canal establishment, similar to our TCDD exposed embryos, include the *dog-earded, valentino, lauscher*, and *colourless* mutant. The *dog-eared* mutant, which disrupts function of *eyes absent-1 (eya1*), has an altered semicircular canal phenotype with defects in maculae and cristae establishment (Kozlowski, Whitfield, Hukriede, Lam, & Weinberg, 2005). The *valentino* mutant displays varying degrees of severity in semicircular canal establishment with some embryos showing partial pillar fusion, which was also observed following TCDD-exposure (Moens, Cordes, Giorgianni, Barsh, & Kimmel, 1998). The *lauscher* mutation caused by a disruption to *gpr124*, an adhesion class G protein-coupled receptor gene, results in defective pillar budding and fusion (Geng et al., 2013). A large deletion mutant b971, which encompasses *sox9b*, produces smaller and misshapen ears and *sox9a* and *sox9b* double mutants lack ears altogether(Yan et al., 2005). Finally, the *colourless* mutation caused by loss of *sox10* results in small ears, abnormal sensory patches, underdeveloped semicircular canals (Dutton et al., 2009; Lopes et al., 2001). In our study, we found that TCDD exposure targets Sox10+ epithelial structures in the ear and that TCDD induced disruption of these structures results in defects in sensory canal establishment. TCDD exposure has also been shown to reduce the number of Sox10+ oligodendrocytes in the developing brain (Martin et al., 2022). Given that in certain developmental contexts *sox10* is downstream of *sox9b*, a known target of the AHR signaling pathway, it could be that TCDD-induced AHR activation indirectly reduces Sox10 expression by impairing Sox9b expression (Yan et al., 2005). Alternatively, as we discuss below, multiple SoxE genes could be direct targets of TCDD-induced toxicity.

### 4.3 SoxE genes as targets of TCDD-induced toxicity

SoxE proteins are a family of transcription factors grouped together by sequence identity in their high-mobility group (HMG) domains (Chiang et al., 2001). The SoxE family in zebrafish is composed of the transcription factors Sox8, Sox9a, Sox9b, and Sox10, which are critical for embryonic development of the otic vesicle, jaw, heart, fin, gonads, CNS, and PNS (Hofsteen et al., 2013; She & Yang, 2017; Stolt & Wegner, 2010; Yan et al., 2005). Previous work has shown that TCDD exposure reduces expression of Sox9b in the developing jaw, heart, and fin (Andreasen, Mathew, & Tanguay, 2006; Burns et al., 2015; Carney et al., 2006; Plavicki et al., 2013; Xiong et al., 2008). Zebrafish exposed to TCDD at 96 hpf show reduced expression of *sox9b* in the jaw at 4 and 12 hours post-exposure (Xiong et al., 2008). In the developing heart, loss of Sox9b can account for a significant subset of the cardiac phenotypes produced by embryonic TCDD exposure (Hofsteen et al., 2013). However, it remained unknown whether other SoxE genes such as *sox9a, sox10*, and *sox8* are TCDD targets.

The otic placode and the developing otic vesicle express Sox9a, Sox9b and Sox10 (Chiang et al., 2001; Dutton et al., 2009) *sox10* expression initially overlaps with *sox9a* and *sox9b*, but eventually expression of *sox10* is maintained in all otic epithelial cells while *sox9a* and *sox9b* expression becomes restricted to specific domains (Dutton et al., 2009). Previous studies have shown that loss of Sox10 results in impaired in otic placode establishment and, ultimately, in structural and sensory deficiencies including defects in the semicircular canals (Dutton et al., 2009). Our data indicate that TCDD targets Sox10+ otic epithelium and that TCDD toxicity results in structural defects.

Future work should be conducted to determine if other SoxE genes are targets of TCDD independent of Sox9b and to understand the downstream targets of Sox10 in the ear. Potential targets include those found in loss of *sox10* function studies. For instance, a loss of function *sox10* mutation in zebrafish results in altered expression of signaling molecules *fibroblast growth factor 8a (fgf8*) and *bone morphogenetic protein 4 (bmp4*) in the ear (Dutton et al., 2009). Fgf8 is required to induce the otic vesicle placode and to form the otic epithelium (Liu et al., 2003).

Correspondingly, in mice, loss of Bmp4 impaired cristae and semicircular canal establishment (Chang et al., 2008) and in zebrafish, excess Bmp4 treatment resulted in ectopic or absent pillar formation, suggesting Bmp4 is involved in semicircular canal establishment. Although previous studies have not investigated the effects of TCDD exposure on expression of Bmp4 and Fgf8 specifically in the ear, previous research found that following TCDD exposure Bmp4 expression was expanded in the heart and, based on whole embryo qPCR data, is globally upregulated in embryos and larvae (Mehta, Peterson, & Heideman, 2008; Yue, Martin, Martin, Taylor, & Plavicki, 2021). TCDD was shown not affect Fgf8 expression following fin amputation (Andreasen, Mathew, Löhr, Hasson, & Tanguay, 2007); however, it is not clear whether Fgf8 expression is altered in other tissues in response to TCDD exposure. Expression analysis of both Fgf8 and Bmp4 in the ear following TCDD exposure would help to further elucidate the molecular mechanisms mediating the phenotypes described in this paper.

### 4.4 TCDD induced dysregulation of collagen deposition

Our studies revealed that TCDD exposure reduces expression of Col2a1, a critical collagen subtype found in many structures including the inner ear. Mutations in COL2A1 can lead to hearing loss in humans (Acke, Dhooge, Malfait, & De Leenheer, 2012). Zebrafish have two *Col2a1* homologues, *col2a1a* and *col2a1b*, with distinct and overlapping expression domains (Dale & Topczewski, 2011). The Col2a1 antibody utilized in this study broadly marks collagen type II and it is therefore not possible for us to distinguish if there are differential effects on the homologues following TCDD exposure. Notwithstanding, we were able to capture a noteworthy reduction of Col2a1 in the otic vesicles suggesting that one or both collagen type II homologs are affected by TCDD exposure. TCDD exposure has also been shown to impair osteogenesis and affect type II collagen expression in medaka; thus indicating that TCDD exposure affects collagen deposition in multiple fish species (Dong, Hinton, & Kullman, 2012; Watson et al., 2017).

In the mouse, *Col2a1* is known to be a direct target of Sox9 (Bell et al., 1997). The observed reduction in Col2a1 is likely the result of inhibited Sox9b expression. The previous studies have shown AHR activation represses *sox9b* function through the expression of a long noncoding RNA, *sox9b long intergenic noncoding RNA (slincR) (slincR)*(Garcia et al., 2018). It is not clear if repression of Sox9b is the primary mediator of the observed otic vesicle phenotypes or, if so, if Sox9b repression is occurring through the same mechanism. Sox9 is also known to be a target of retinoic acid signaling (Dale & Topczewski, 2011; Williams, 2022). A previous study found that retinoic acid signaling suppressed SOX9, the human orthologue of zebrafish Sox9b, in chondrocytes and reduced *Col2a1 e*xpression (He, Brysk, Tyring, Ohkubo, & Brysk, 2001). Conversely, overexpression of SOX9 reversed retinoic acid induced suppression of *Col2a1* (He et al., 2001). Similar regulatory interactions between Sox9 and retinoic acid have been observed in the testis (Bowles et al., 2018). These studies indicate that retinoic acid is an important regulator of *Sox9* and *Col2a1* expression.

Retinoic acid signaling has also been implicated in zebrafish ear development, specifically regulating the development of the semicircular canals and otoliths. Disruption of this pathway results in a circling behavior previously associated with inner ear malformations (Mackowetzky et al., 2022). A number of lines of evidence suggest an interaction between TCDD exposure and retinoic acid signaling in the generation of adverse outcome (Fiorella, P. D., Olson, J. R., & Napoli, 1995; Herlin et al., 2021; Jacobs et al., 2011; Murphy, Villano, Dorn, & White, 2004; Schmidt et al., 2003; Weston, W. M., Nugent, P., & Greene, 1995). Consequently, the retinoic acid pathway presents exciting prospects for future studies of TCDD-induced disruption of otic vesicle development.

## 5 Conclusions

This study reveals the otic vesicle as a novel target of embryonic TCDD toxicity. Through this work, we characterized the auditory-vestibular structures and developmental processes in otic vesicle formation affected by early TCDD exposure. Specifically, our data demonstrates that embryonic exposure to TCDD leads to defects in pillar establishment, pillar fusion, and topography, ultimately leading to impaired development of the ear’s semicircular canals required for detecting angular rotation. Furthermore, we found TCDD significantly inhibited expression of collagen type II, a collagen subtype with important functions in mammalian ear development. Future studies will help determine the molecular mechanisms that produce otic phenotypes following TCDD exposure.

## 7 Author Contributions

L.G.C-R and J.S.P. designed the research study, developed figures, and wrote the manuscript. L.G.C-R performed TCDD exposures, collected, and prepared samples. L.G.C-R, R.P., and R.C. performed confocal imaging and acquired data for analysis. L.G.C-R, G.O., A.P., and N.B. conducted data analysis. L.G.C-R is supported by the National Science Foundation Graduate Research Fellowship (NSF-GRFP and previously by the NIEHS Training in Environmental Pathology T32 (ES007272). J.S.P. is supported by an NIEHS K99/R00 (ES023848), a CPVB Phase II COBRE (2PG20GM103652), and an NIEHS ONES award (ES030109).

## 8 Acknowledgements

The authors express deep appreciation for all the past and present members of the Plavicki Lab. They would like to thank all members for their help in maintaining the lab and for their feedback. In particular, the authors would like to thank Shannon Paquette for her valuable input. Lastly, the Collagen Type II (II-II6B3) developed by T.F. Linsenmayer was obtained from the Developmental Studies Hybridoma Bank, created by the NICHD of the NIH and maintained at The University of Iowa, Department of Biology, Iowa City, IA 52242.

