## Supplemental Figures for "Exposure to the persistent organic pollutant 2,3,7,8-Tetrachlorodibenzo-p-dioxin (TCDD, dioxin) disrupts development of the zebrafish inner ear"

**10 Supplemental Figures**

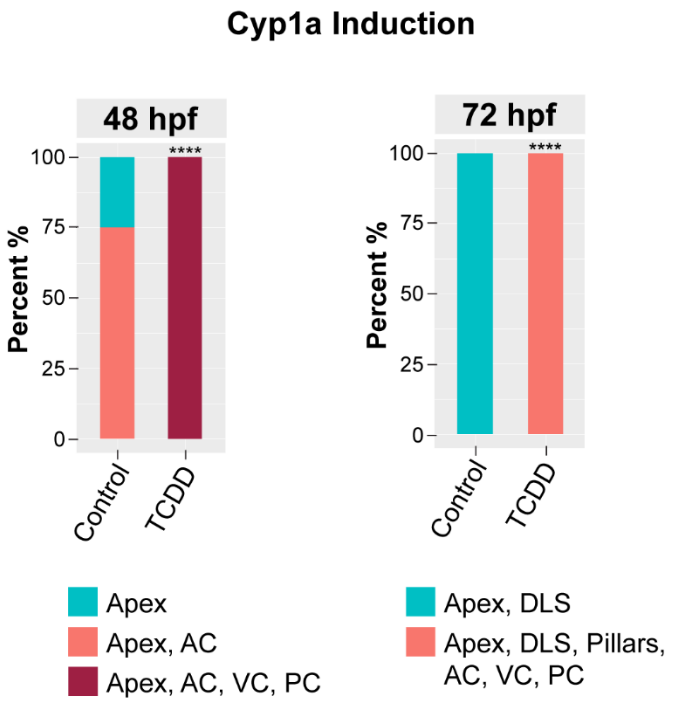


**Supplemental Figure 1. Quantification of Cyp1a induction in the developing zebrafish ear.** Percentage of ear structure with Cyp1a induction in control and TCDD-exposed embryos at 48 and 72 hpf. Abbreviations: anterior crista (AC), ventral crista (VC), posterior crista (PC). n=8-10/group (48hpf) and n=10-13/group (72hpf) with a minimum of 3 replicates. ****p < 0.0001

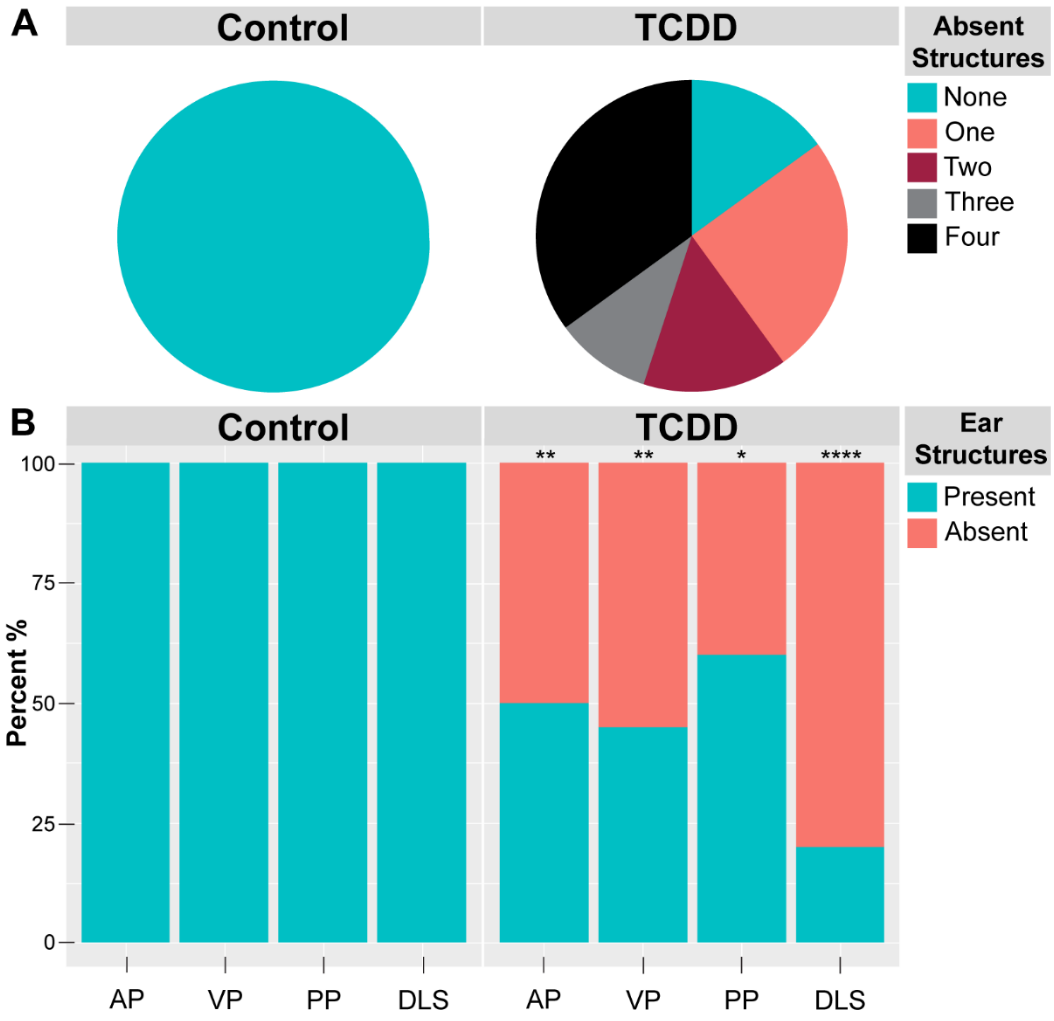


**Supplemental Figure 2. Quantification of structural deficits at 96 hpf.** Percent of coincident absent structures in control and TCDD exposed embryos at 96 hpf (A). Percent of ear structure absences (B). Abbreviations: anterior pillar (AP), ventral pillar (VP), posterior pillar (PP), dorsolateral septum (DLS). n=12-20/group with a minimum of 3 replicates. *$p < 0.05$, **$p < 0.01$, ***$p < 0.001$, ****$p < 0.0001$.

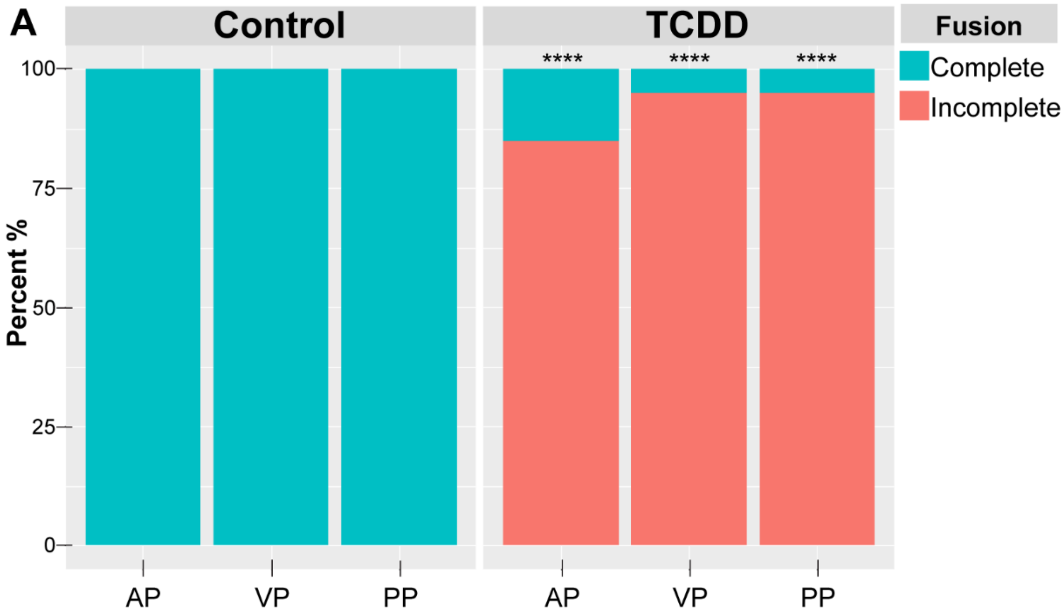


**Supplemental Figure 3. Incomplete pillar fusion quantification at 96 hpf.** Percent of structures that failed to establish a connection at fusion plate in control and TCDD exposed embryos at 96 hpf (A). Abbreviations: anterior pillar (AP), ventral pillar (VP), posterior pillar (PP). n=12-20/group with a minimum of 3 replicates. ****$p < 0.0001$

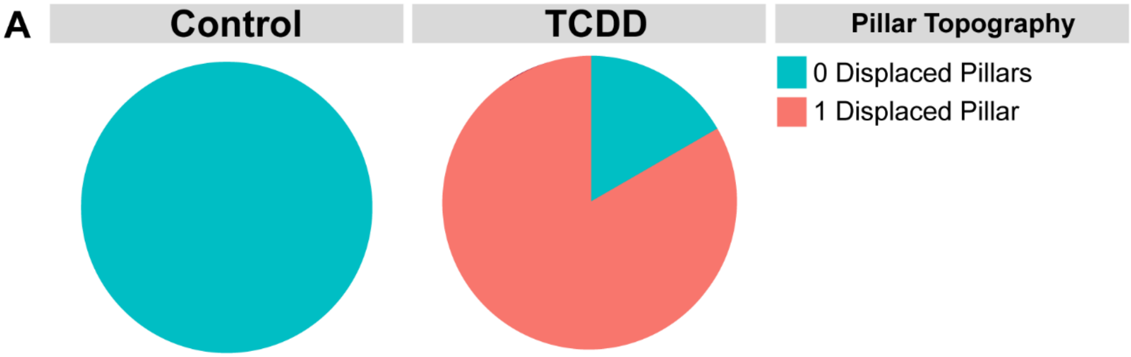


**Supplemental Figure 4. 96hpf pillar topography defect quantification.** Percent of control and TCDD exposed embryos with zero or one displaced pillars at 96 hpf hpf (A). n=12-20/group with a minimum of 3 replicates.

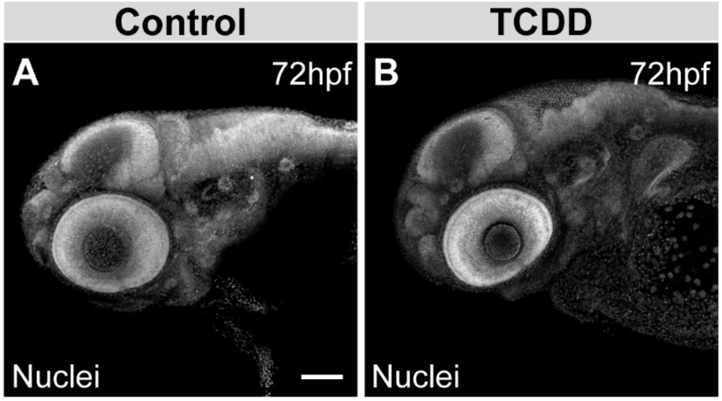


**Supplemental Figure 5. Eye morphology.** Control (A) and TCDD exposed (B) embryos at 72 hpf stained for DAPI showing different eye morphology phenotypes.

**Supplemental Movie 1. Time-lapse microscopy of pillar budding and fusion in the ear** Control (Video 1) and TCDD exposed (Video 2) ears developing *in vivo* from 52 hpf – 72 hpf; video captured through confocal microscopy at 10x magnification. n=5/group with 3 replicates. Abbreviation: otic vesicle (OV)
